# L-DOPA Rescues Reach Height but Not Trajectory Area in a Bimanual String-Pulling Behavior in Parkinsonian Rats

**DOI:** 10.64898/2026.10.26.761430

**Authors:** Matthew B Schmit, Asier Aristieta, Rodica Caparu, Aryan J Malik, Audrey White, Yana Surtchev, Vivianna Pederson, Mitchell J Bartlett, Abhilasha Vishwanath, Torsten Falk, Stephen L Cowen

## Abstract

While gross motor changes in Parkinson’s disease, bradykinesia, hypokinesia, difficulty initiating movement, have been well studied in rodent models of the disease, limb movements, and especially bimanual coordination deficits, are more difficult to study in these animal models. Here we describe a novel bimanual string-pulling task that can be used to generate a rich repertoire of behaviors that require limb and bimanual coordination in rats with unilateral 6-hydroxydopamine (6-OHDA) lesion. We show that 6-OHDA lesion significantly reduces string speed, pulling bout duration, and paw speed especially during reaching, and reduces the area of paw trajectories during pulling. However, it does not affect the variability around that movement trajectory. We also show that levodopa (L-DOPA) treatment restores many of these measures but not all. L-DOPA restores paw speed and string-pulling speed and fully restores paw reach height and pulling bout duration but has no effect on trajectory area. Together, our results demonstrate that dopamine depletion differentially effects features of this bimanual coordination task, that L-DOPA mostly, but not fully, restores string pulling performance, and opens up a new avenue for measuring the effect of Parkinson’s disease treatments on limb movement, allowing for better assessment of how effectively treatments restore different types of motor activity.

## Introduction

Some of the debilitating motor symptoms of Parkinson’s diseases include bradykinesia^1,2^, hypokinesia^3^, loss of motor control^4,5^, difficulty manipulating small objects^6^, and problems with bimanual coordination^7,8^. These symptoms emerge following the progressive loss of dopaminergic neurons in the substantia nigra pars compacta. While gross motor symptoms can be studied effectively in rodents, studying limb movement and especially bimanual coordination is more difficult. In rats, limb motor performance is often studied using tasks such as the vermicelli handling task^9^, lever pressing tasks^10,11^, the staircase test^12^, and gait analysis^13–16^, but these tasks generally require intense manual analysis and most to not assess limb trajectory or bimanual coordination.

A rich dataset is key for teasing apart which aspects of motor movements are affected by disease and treatment. For example, dopamine depletion and striatal ensemble silencing have strong effects on the speed and vigor of limb movements, but less effects on movement trajectories, suggesting these two are dissociable^11^. On the treatment side, while stride length and arm swing range of motion are improved by levodopa (L-DOPA), the most common drug prescribed for Parkinson’s disease, postural measures and movement initiation are more resistant to pharmacological treatment^17,18^. L-DOPA treatment does not always restore motor function uniformly. For example, finger tapping, a common diagnostic of Parkinson’s disease, was not significantly improved by L-DOPA treatment and tapping rate even worsened in patients without previous treatment^19^, although this is not consistent across patients who were already taking L-DOPA^20^.

Detailed motor analysis of inter-limb movement can help us better understand how dopamine depletion and pharmacological interventions affect motor performance. Here, we use a novel bimanual string pulling task^21,22^ to asses coordinated motor behaviors in a 6 hydroxydopamine (6-OHDA) rat model of Parkinson’s disease^23,24^ before and after L-DOPA treatment. We used the 6-OHDA model as it is a standard model of PD, is a robust model of dopamine depletion, and permits the use of the contralateral paw as a control.

String pulling requires precise bimanual forelimb coordination, and has been used to assess motor behavior^25^ across many species, including humans^26,27^. Recently, it has been used in a rat model to investigate movement disorders resulting from stroke^28^. Using a high speed camera and markerless tracking^22,29^, the task allows precise tracking of paw position, allowing the investigation of individual paw trajectories, speeds, movement variability, changes in specific phases of pulling (e.g., reaching upwards vs. pushing downwards on the string^21^) and relationships between the paws. In addition, since weaker grip strength is a common symptom of Parkinson’s disease^3^, we included a measure of string tension to determine the amount of force exerted by each paw during pulling.

Compared to pre-lesion levels, 6-OHDA lesion significantly reduced total string pulled and string-pulling speed, as well as average bout duration. We also show that, as seen in micrographia^30^ and hypokinesia^3^, paw trajectories during pulling were significantly smaller. We show that paw speed is significantly slower when compared to the intact paw across both the reaching and downward phases of pulling, but that reaching is especially affected by the 6-OHDA lesion. In agreement with this observation, we show that the tension placed on the string during pulling does not differ between lesioned and unlesioned animals. Finally, we show that L-DOPA treatment, while improving many measures of string pulling, is most effective at restoring bout duration and reach height.

## Methods

### Animals

Adult male Sprague-Dawley rats (n = 12 6-OHDA lesion, 5 sham lesion; ∼250 g upon arrival, Envigo RMC Inc., Indianapolis, IN) were single housed in a temperature and humidity-controlled room on a 12-hour reverse light/dark cycle (lights off between 9 a.m. and 9 p.m.). Food and water were provided *ad libitum* during the habituation and surgical recovery periods. Prior to training, each animals’ initial body weight was measured and recorded. During behavioral training, to maintain task motivation, animals were food restricted to maintain 85% initial body weight measured following each behavioral session. All animal studies were approved by the University of Arizona Institutional Animal Care and Use Committee and in accordance with the NIH ARRIVE and the Care and Use of Laboratory Animal guidelines.

### Behavioral Training

Prior to behavioral training on the string-pulling task, animals were handled for 30 minutes per day for one week to habituate to handling and room. Animals were trained on an adapted version of the protocol described in Jordan et. al^22^. Food neophobia was mitigated by providing cheerios in the home cage for the first 3 days of habituation. All training sessions lasted 30 minutes. The training protocol consisted of two phases as described below:

#### Phase One

Animals were placed in a box (11 x 11 x 16 in) with a strings of varying length were draped over the edge of the box, each baited with a Cheerio tied to the end. The string was also coated with crushed Cheerio dust to incentivize pulling. The length of the string was progressively increased after animals successfully pulled that length and acquired the Cheerio reward 3 times (5, 10, 30, 45, 60 cm). Animals were allowed to continue pulling the 60 cm length string for reward until the end of the 30-minute training session. Each animal was required to pull the 60 cm string 3 times on 3 consecutive days to advance to phase two.

#### Phase Two

Animals were placed on the string-pulling platform and trained on a progressive distance protocol where three complete pulls were required at each length before advancing to the next length (5, 10, 20, 50, 100, 150, 200, 250 cm). Each successful pull was rewarded with a Cheerio. Once the full protocol was completed for a session (completed up to 250 cm 3 times), animals were allowed to pull additional bouts at the 250 cm length until session completion at 30 minutes. The system automatically tracked distance pulled as previously described^22^. Reward and distance expectations were reinforced by an audible click provided by the system when the required distance was pulled for each step. Pre-lesion, each animal was required to perform three consecutive days to protocol completion as a performance baseline.

All animals completed the training and met the required performance baselines within two weeks. Post lesion, rats were retrained in the same way as untrained rats until they had received 3 weeks of training.

### Stereotaxic Surgery

6-OHDA lesions were performed as previously described^31^. Rats were inducted with 5% isoflurane (VetOne, Boise, ID) and kept between 1-2% during surgery. Animals were injected with the antibiotic Gentamicin (8 mg/kg, i.p.; VetOne) and analgesic Ketoprofen (5 mg/kg, i.p.; Zoetis) prior to surgery. Desipramine (12.5 mg/kg, i.p.; Sigma-Aldrich, dissolved in 10% dimethyl sulfoxide with 0.9% sterile saline) was administered 20 minutes prior to 6-OHDA injection to protect noradrenergic cells. 6-OHDA (Sigma-Aldrich) was dissolved in 0.02% L-ascorbic acid vehicle (Sigma-Aldrich; in 0.9% sterile saline, VetOne, 5mg/ml). 6-OHDA was injected (2.5 µl/site at 0.3 µl/min) with a microinjector (Stoelting Co., Wood Dale, IL) connected to a needle (Gastight #1701, Hamilton Co., Reno, NV) targeting the medial forebrain bundle (MFB) at two coordinates (AP −1.8; ML +2.0; DV −8.2 mm and AP −2.8; ML +1.8; DV −8.2 mm, with all DV measurements from brain surface). Sham-treated animals were injected with 0.02% L-ascorbic acid vehicle at the same two coordinates. Needles were left in place for 5 minutes following completion of the injection to prevent backflow upon retraction.

### Drug Formulation

Levodopa (L-DOPA; 6 mg/kg, i.p.; Sigma-Aldrich) was formulated in 0.9% sterile saline (VetOne) with 12 mg/kg benserazide (Sigma-Aldrich). Control injections consisted of 0.9% sterile saline vehicle (VetOne). All injections were given 20 minutes prior to each string-pulling session.

### Near-Infrared Tyrosine Hydroxylase Immunohistochemistry

6-OHDA lesions and intact sham lesion characterization was done with near-infrared immunohistochemistry for tyrosine hydroxylase (TH). At the experimental endpoint, animals were deeply anesthetized with Euthasol pentobarbital solution (85.8ml/kg VetOne, i.p.) and transcardially perfused with phosphate-buffered saline then 4% formaldehyde solution. Brains were extracted and sectioned with a cryostat (CM1850, Leica, Wetzlar, Germany). Three coronal sections from AP, +1.2 mm; AP, +0.48 mm; AP, −0.26 mm relative to bregma were selected for each animal according to a rat brain atlas (Paxinos & Watson, 2007). Sections were washed 4 x 5 minutes in Bio-Rad Tris-buffered saline (TBS) blocking buffer (Bio-Rad Laboratories, Hercules, CA) + 0.1% Tween-20 (TBST; Sigma-Aldrich) and then incubated in Liberate Antibody Binding® (L.A.B.; Polysciences, Warrington, PA) solution for 15 minutes to free antibody binding sites following fixation. Sections were then washed 2 x 5 minutes in TBST and incubated in TBS blocking buffer (10% donkey serum (Sigma-Aldrich), 0.5% Triton X-100 (Sigma-Aldrich) and 0.3 M Glycine (Fisher BioReagents, Ottawa, ON)) for 2 hours at room temperature. Sections were then incubated in Intercept® TBS blocking buffer (LI-COR Biosciences, Lincoln, NE) with 1% donkey serum, 0.2% Triton X-100, and rabbit anti-TH (1:4000; Millipore, St. Louis, MO) overnight at 4°C. Following overnight incubation in primary antibody, tissue was washed 4 x 5 minutes in TBST and incubated in TBS blocking buffer (1% donkey serum, 0.2% Triton X-100) and IRDye® 800CW Donkey anti-Rabbit (1:200; LI-COR Biosciences) for 2 hours at room temperature shielded from light. Tissue was then washed 4 x 5 minutes in TBST and plated on charged slides (Fisherbrand, Pittsburgh, PA) with VECTASHIELD® HardSet Antifade Mounting Medium (Vector Laboratories, Newark, CA). Prepared slides were imaged with an Odyssey M scanner (LI-COR Biosciences). TH-immunoreactivity was quantified using Image Studio 6.2 software (LI-COR Biosciences). Mean signal intensity was measured in the striatum for each hemisphere, and background subtracted for non-specific antibody binding using signal measured in the corpus callosum^31,32^

### String-Pulling Apparatus

The string-pulling system consists of a loop of string mounted on pulleys with one pulley connected to a digital rotary encoder (BQLZR, Shenzhen, China) to measure string speed and distance pulled, described below. The signal was monitored by a connected Arduino microcontroller, which also controlled a solenoid used as an audio cue for reward delivery. Using 3 extra rollers in the string path, with one mounted to a load sensor, we measured tension in the string (See **Figure 6b**). Details regarding construction can be found in Jordan et al.^22^. Signals from the rotary encoder and load cell were acquired using a NI PXIe-1071 (National Instruments Inc.,) that was controlled by the SpikeGLX software package (Bill Karsh, Janelia Research Campus, https://billkarsh.github.io/SpikeGLX).

### Video Acquisition and Paw Tracking

A front-facing camera (MakoU130b, Allied Vision, Stadtroda, Germany) monitored string-pulling. Data was acquired at 330 fps using StreamPix software (NorPix, Montreal, Quebec). Forelimb and nose position were extracted from the video using DeepLabCut^22^. Custom MATLAB code was used for further analysis and provided at https://github.com/CowenLab/String_Pull_PD_334.)

### String Speed Analysis

The rotary encoder produced a pulse for every 1 cm of string pulled and this signal was used to calculate the following values: ***String Speed:*** Average string speed during good pulling periods, defined as string speed > 1 cm/s. ***Mean Peak Speed:*** Average of peak string speeds (calculated using findpeaks, MATLAB) during good pulling periods. ***Interpeak Interval:*** Median distance between string speed peaks after rejecting inter-peak intervals > 1 s. ***Bout Duration:*** The mean duration of good pulling periods.

### String Tension Analysis

Our string tension meter used pulse width modulation (10 ms pulse width) to encode relative string tension from a pre-session baseline (Github link for Audrey PWM code). Average string tension for a session was defined as pulse width divided by 10ms, averaged over good pulling periods.

### Paw Position Processing

To process paw X and Y position from DeepLabCut we first we rejected all points with a likelihood measure of less than 0.7. Then we applied a moving median with a window size of 4 points (12 ms, data sampled at 330 Hz) and a moving mean with a window size of 8 points (24 ms), ignoring rejected values. We then linearly interpolated across gaps in the data less than 20 points (60 ms). Finally, we smoothed our cleaned data with a Savitzky-Golay filter of order 7 and window size of 25 points (75 ms), smoothing out noise while preserving sharp changes in position expected in string-pulling. From these values we calculated paw position and converted to centimeters.

### Paw Trajectory Measures (In Figure Order)

To analyze how a paw moves while repeatedly pulling the string, we first needed to segment behavior into individual pulls and then process these segmented pulls to compare their shapes across different durations and positions. We first defined a pull cycle as the time between two paw Y maxima (defined using findpeaks, MATLAB). Pull cycles longer than 1 s were rejected as were the resulting bottom and top 10% of cycles. To average trajectories across pulls of different durations we interpolated both X and Y position at 100 points from cycle start to end. We then subtracted the mean X and Y coordinates from each cycle’s trajectory and averaged these centered, interpolated, trajectories across all pull cycles.

***Trajectory Area*** Trajectory area was defined as the area inside the centered and averaged paw trajectory across pull cycles. ***Reach Y Range:*** Defined as the difference between the highest and lowest Y points of the pull cycle trajectories, averaged across cycles. ***Reach Height***: Defined as the maximum Y coordinate averaged across pre-centering pull-cycle trajectories. ***Average Path Departure:*** Defined as the mean Euclidian distance of each individual centered and interpolated cycle trajectory from the average trajectory that session.

### Trajectory Independent Measures

***Time to Next Pull:*** First we identified the start of each pull for a single paw. We kept only pulls where the opposite paw pulled and then the original paw pulled again, eliminating pulls from the start and stop of a pulling bout. We then calculated the time from these cleaned pulls to the opposite paw’s next pull. ***Paw Speed:*** Calculated as the average paw speed across a trace during good pull periods, defined as where both the left and right paws were tracked and the duration of the good data segment was greater than 1 s. ***Paw Speed During Reach and Downward Phases:*** Paw speed was defined as the average speed during reach and downward phases: the reach phase of pulling was defined as contiguous periods where the paw Y velocity was positive, the distance traversed in that period was at least 1.5 cm, and the paw remained above 5 cm while the downward phase was defined as the velocity being negative, the distance traversed at least 1.5 cm, and the height of the paw above 5 cm.

### Statistics

Prior to performing hypothesis tests, a Kolmogorov-Smirnoff test for normality determined whether a parametric or non-parametric hypothesis tests was subsequently used. Effect size was indicated as Cohen’s d for when parametric tests were applied, and the r measure of effect size was used following non-parametric tests^33,34^.

For comparisons across multiple conditions, we used a repeated-measures ANOVA (RMANOVA), first running a sphericity test to determine if a Greenhouse-Geisler correction was necessary or not. Then, if the test was significant, we ran multiple paired tests on the comparisons we determined were relevant prior to running the RMANOVA and applied a Bonferroni-Holm correction to the p-values.

## Results

### Pre-training and Pre-lesion Performance on the String-pulling Task

In order to study how a unilateral 6-OHDA lesion and L-DOPA treatment affect motor skills, we first trained rats on a string-pulling paradigm as described previously^22^ (**Figure 1a**). Prior to lesion, rats pulled an average of 8,770 cm (SEM ±798 cm) of string per session and averaged 25.99 cm/s (SEM ±0.89 cm/s) during pulling bouts. **Figure 1b** shows string speed during a single pulling bout. There was clear rhythmicity in string speed, with the peaks indicating increased speed due to the downstroke of each paw. An autocorrelogram of string speed showed the first peak in rhythmicity occurring at 6 Hz (**Figure 1c**).

**Figure 1:**
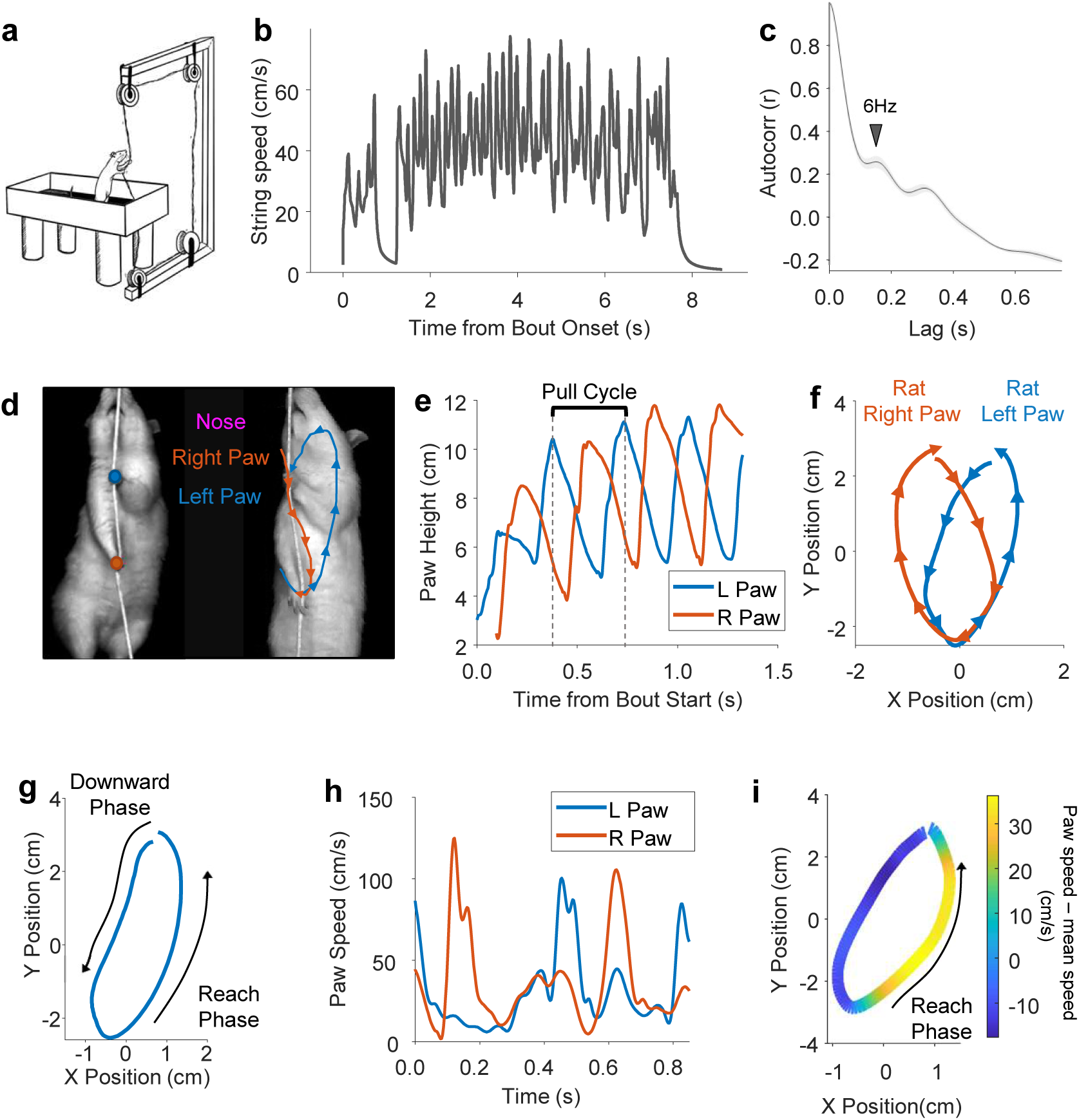
a. Schematic of string-pulling set up, with rat on platform pulling the string with both paws. b. Example string speed trace from one pulling bout c. Autocorrelogram of string speed showing a first peak at 6 Hz. d. Example image showing paw tracking (left) and paw trajectories over time (right). e. Example trace of left and right paw height during the start of a pulling bout. A pull cycle was defined as starting at the peak of that paw’s height until the next peak. f. Averaging the trajectories of each paw across pull cycles we can produce an average movement trajectory per paw. String would be in the center of this diagram. g. Each pull cycle can be divided into a downward phase and an upward ‘reach’ phase. h. Example paw speed traces showing peaks in speed during pulling. i. Paw trajectory with paw speed overlayed in color, showing an increase in paw speed during reaching.

To compare the motion of left and right paws, marker-less tracking via DeepLabCut^29^ was used to track individual paw movement during pulling (**Figure 1d**). The height of each paw peaked at the start of its respective pull cycle (**Figure 1e)**, with an average of 800 cycles (SEM ±45 cycles) per paw per session. The average trajectory of each paw was computed by first mean-centering each pull trajectory and then averaging across all cycles (**Figure 1f**, see Methods). Trajectories were divided into two behavioral segments, an upward reach phase, where the rat releases the string and reaches upward, and a downward phase where the rat grasps the string and moves it downward, combining the push and pull phases defined in prior work^21^ (**Figure 1g**). Each paw shows periods of higher speed corresponding to the reach (**Figure 1h**). Taking the average speed during a pull cycle and projecting it onto a pull trajectory we can see that the paw speed peaks during the upward reaching phase (**Figure 1i**). Together, these data illustrate that string pulling contains a rich repertoire of motor measures, both bimanual and performance based, that may be impacted by Parkinson’s disease.

### 6-OHDA Lesion Reduces String-Pulling Speed and Length Pulled

To investigate how motor skills are affected by Parkinson’s disease, we injected either 6-OHDA or vehicle into the right medial forebrain bundles of two cohorts of rats to preferentially ablate nigral dopaminergic neurons and striatal terminals of one hemisphere. Using immunohistochemistry, tyrosine hydroxylase (TH), a marker of dopaminergic neurons and terminals, was significantly depleted in the striatum of 6-OHDA but not the sham group (**Figure 2a-b**, unpaired t-test, n = 5 intact 11 lesioned, t = 51.8, p = 2.12e-17, Cohen’s d = 26.4).

**Figure 2:**
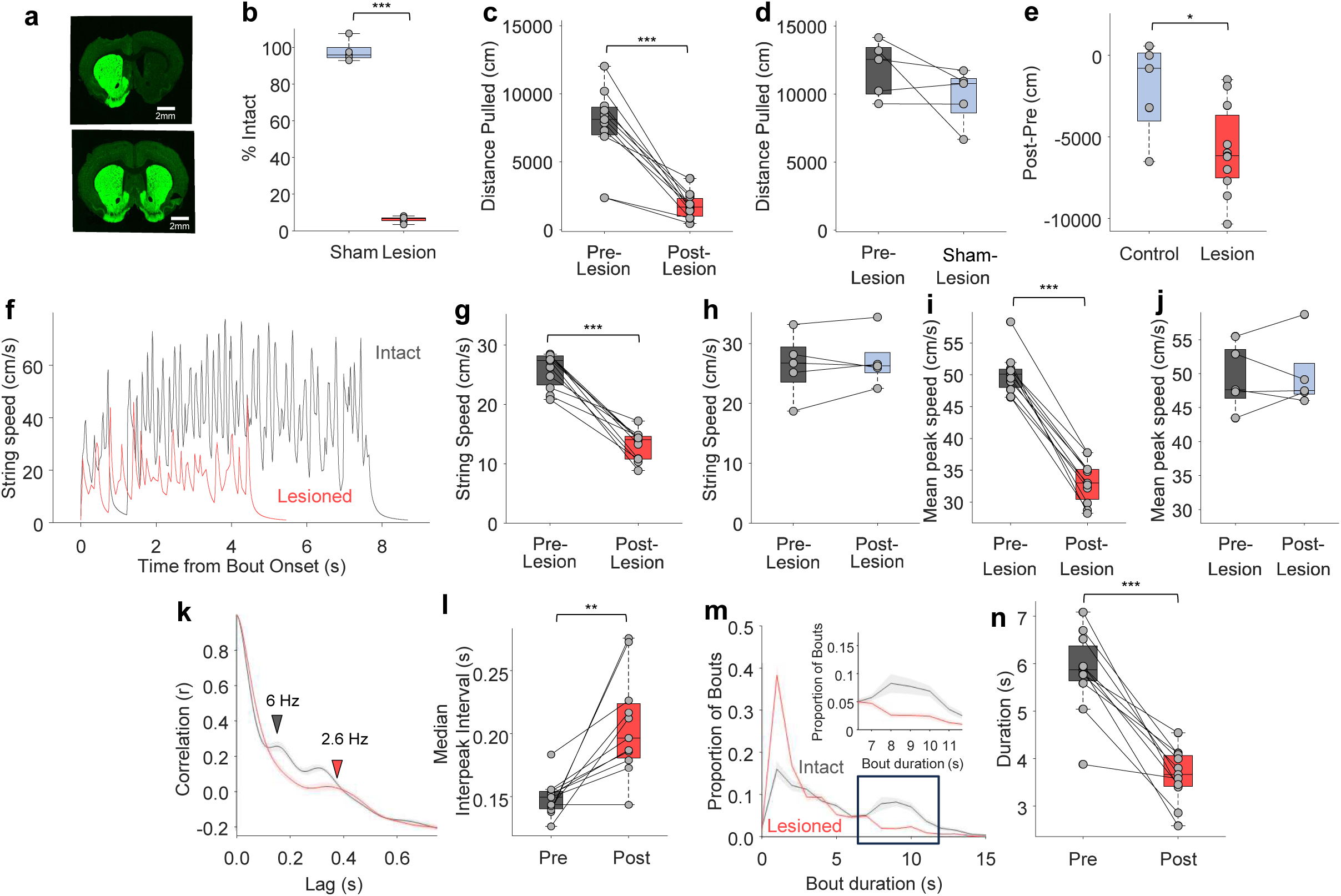
a. Representative histology from 6-OHDA lesion and sham-lesion animals showing TH (green) levels reduced in lesioned rats. b. 6-OHDA lesion significantly reduced TH staining on the lesioned side vs sham (unpaired t-test n = 11 6-OHDA, n = 5 sham-lesion rats, p = 2.12e-17). c. Distance pulled pre- and post-lesion was significantly reduced (paired t-test, n = 11, p = 3.56e-05). d. Distance pulled pre- and post-sham-lesion showed no difference (paired t-test, n = 5, p = 0.198). e. Lesioned rats pulled significantly less distance than sham-lesioned rats(t-test, n = 11 6-OHDA, n = 5 sham-lesion rats, p = 0.0238 f. Example traces of string speed during a pulling bout pre- and post-lesion. g. Average string speed during a pulling bout was significantly lower post-lesion vs pre (paired t-test, n = 11, p = 1.53e-06). h. String pulling speed was not affected post sham-lesion (paired t-test, n = 5, p = 0.495). i. Peak speed was significantly reduced post-lesion (paired t-test, n = 11, p = 1.27e-06). j. Peak speed was not affected post sham-lesion (paired t-test, n = 5, p = 0.366). k. Autocorrelagram of string speed averaged over 9 mice before and after lesion showing a slower first peak. Shaded area is S.E.M. l. Mean inter-peak interval, computed on string-pulling speed traces, was significantly longer post-lesion (paired t-test, n = 11, p = 0.00179). m. Distribution of bout lengths before and after lesion, showing the proportion of long bouts decreasing as the proportion of short bouts increasing. Shaded area is S.E.M. n. Average pulling bout duration was significantly decreased post-lesion (paired t-test, n = 11, p = 2e-05). * = p < 0.05, ** = p < 0.01, *** = p < 0.001

In human patients, Parkinson’s disease is characterized by slow movement and a reduced capacity to sustain effortful behaviors^1^. Relative to pre-lesion, 6-OHDA lesioned rats pulled significantly less distance per session (**Figure 2c**, paired t-test, n = 11, t = −7.04, p = 3.56e-05, Cohen’s d = −1.96). Sham lesioned rats did not show a significant decrease in total distance pulled following sham lesion (**Figure 2d**, paired t-test, n = 5, t = −1.54, p = 0.198). The 6-OHDA group pulled significantly less than the sham group relative to pre-lesion (**Figure 2e**, t-test, n = 11 6-OHDA, n = 5 sham-lesion rats, t = −2.53, p = 0.0238, Cohen’s d = −1.29). **Figure 2f** presents two examples of pulling bouts illustrating how string-pulling speed and bout duration were reduced post lesion. String speed was measured within each active pulling bout where bouts were defined as periods when string speed exceeded 1 cm/s. We found that mean string speed during pulling was significantly slower post 6-OHDA lesion vs pre (**Figure 2g**, paired t-test, n = 11, t = 10, p = 1.53e-06, Cohen’s d = 2.79), while the sham lesion group showed no difference (**Figure 2h**, paired t-test, n = 5, t = −0.75, p = 0.495). Consistent with the reduction in mean speed, the peak speed per cycle was also significantly reduced in the 6-OHDA animals (**Figure 2i**, paired t-test, n = 11, t = 10.2, p = 1.27e-06, Cohen’s d = 2.85) while in shams it was unchanged (**Figure 2j**, paired t-test, n = 5, t = −1.02, p = 0.366).

Further, the rhythmicity of the pulling behavior was disrupted in the 6-OHDA group as indicated by the autocorrelogram of string speed during pulling bouts (**Figure 2k**). Specifically, the inter-peak interval, defined as the median time between string speed peaks, was significantly longer post lesion (**Figure 2l**, paired t-test, n = 11, t = −4.21, p = 0.00179, Cohen’s d = −1.17). The duration of each continuous pulling bout was also measured. Examination of the histogram of bout durations indicated that lesioned animals exhibited a reduction in long pulling bouts with durations of > 5 s (**Figure 2m**,see box).

Indeed, the average duration of bouts was significantly reduced post lesion (**Figure 2n**, paired t-test, n = 11, t = 7.53, p = 2e-05, Cohen’s d = 2.09). Overall, measures of string-pulling performance such as total amount of string pulled, string speed, and pulling bout duration were reduced in lesioned animals, consistent with reduced movement speed and motor effort in individuals with Parkinson’s disease.

### High Speed Paw Tracking Reveals Smaller Movement Trajectories in 6-OHDA Animals

One of the strengths of the 6-OHDA model is its asymmetry, where one paw is dopamine depleted the other is relatively intact^15,23^. Here, we leverage this asymmetry by using DeepLabCut to track each paw before and after 6-OHDA lesion. Post lesion, the average trajectory of the lesioned paw becomes smaller and thinner (**Figure 3a**). The trajectory area was significantly smaller in the lesioned vs intact paw post lesion (**Figure 3b**, Pre-lesion: paired t-test, n = 9, t = 0.465 p = 0.655. Post-lesion: paired t-test, n = 9, t = −6.97, p = 0.000116, Cohen’s d = −2.1). This was not the case after sham lesion (**Supplementary Figure 1a**, Paired t-tests with Bonferroni-Holm correction, n = 5. Pre-lesion: t = −0.461 p = 0.669. Post-lesion: t = −1.74 p = 0.314).

**Figure 3:**
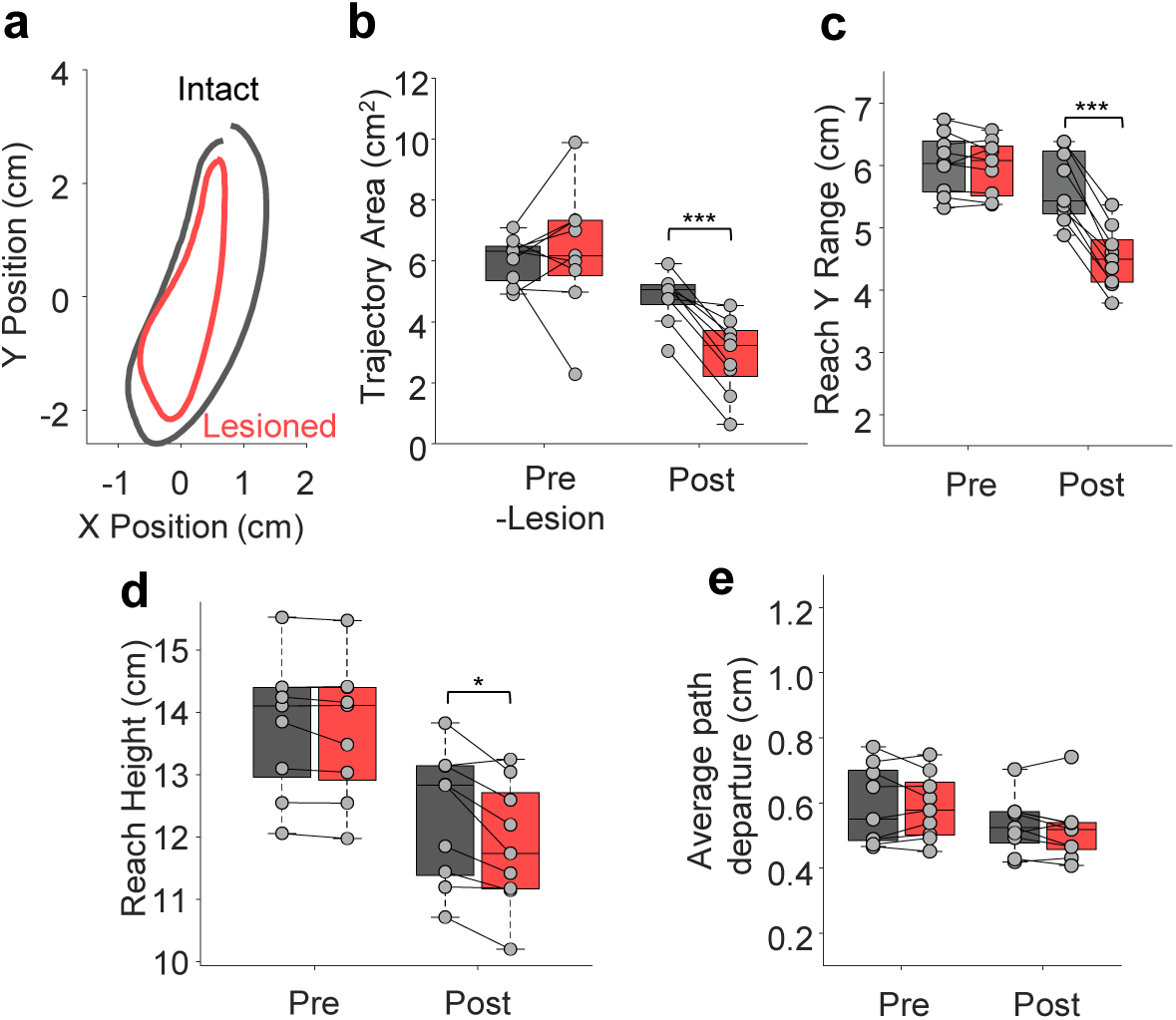
a. Average paw trajectory pre and post 6-ODHA lesion for one rat. b. Post-lesion, paw trajectory area is significantly reduced relative to the non-lesioned paw (paired t-tests with Holm correction, n = 9, Pre-lesion: p = 0.655, post-lesion: p = 1.16e-4) c. The range in the y direction was significantly reduced in the lesioned paw relative to the non-lesioned paw (paired t-tests with Holm correction, n = 9. Pre-lesion: p = 0.391, post-lesion: p = 1.87e-06). d. The absolute height of the reach was significantly lower in the lesioned vs non-lesioned paw (paired t-tests with Holm correction, n = 9. Pre-lesion: p = 0.112, post-lesion: p = 0.0103) e. Average path departure, the average distance per pull from the average path, was not significantly different pre and post-lesion (paired t-tests with Bonferroni-Holm correction, n = 9, pre-lesion, p = 0.842, post-lesion, p = 0.434).

Measuring the Y range of the paw trajectory, we found that the lesioned paw had a significantly smaller Y range compared to the intact paw post lesion (**Figure 3c**, Paired t-tests with Bonferroni-Holm correction, n = 9. Pre-lesion: t = −0.906 p = 0.391. Post-lesion: t = −12.2, p = 1.87e-06, Cohen’s d = −3.68). In sham animals, the sham lesioned paw had a significantly larger Y range both pre and post lesion (**Supplementary Figure 1b**, Paired t-tests with Bonferroni-Holm correction, n = 5. Pre-lesion: t = 3.02, p = 0.0392, Cohen’s d = 1.08. Post-lesion: t = 4.66, p = 0.0192, Cohen’s d = 1.66).

In addition to the Y range of the trajectory, we also measured the absolute height of the reach. We found that the lesioned paw was reaching lower than the intact paw post lesion (**Figure 3d**, paired t-tests with Bonferroni-Holm correction, n = 9. Pre-lesion: t = −1.78 p = 0.112. Post-lesion: t = −3.81, p = 0.0103, Cohen’s d = −1.15), indicating that both the range of movement and its absolute height were affected. In sham animals, both paws had similar heights (**Supplementary Figure 1c**, Paired t-tests with Bonferroni-Holm correction, n = 5. Pre-lesion: t = 1.58, p = 0.189. Post-lesion: t = 2.23 p = 0.179).

Although the trajectory area was reduced post-lesion, the variability around the trajectory was unchanged. Measuring the average path departure, the average distance of each individual pull cycle from the average trajectory, we found it was not significantly different lesion vs intact (**Figure 3e**, paired t-tests with Bonferroni-Holm correction, n = 9.

Pre-lesion: t = 0.206, p = 0.842. Post-lesion: t = 1.34, p = 0.434). In sham animals, we showed a lower average path departure in the lesioned paw (**Supplementary figure 1d**, Paired t-tests with Bonferroni-Holm correction. Pre-lesion: n = 5, t = 2.18, p = 0.095. Post-lesion: t = 3.95, p = 0.0335, Cohen’s d = 1.41). Taken together, we can show that, while the string-pulling trajectories are smaller and lower post-lesion, the variability around that trajectory is relatively low. The pulling behavior remains stereotypical, maintaining its overall shape, while specific characteristics of that shape change post lesion.

### Paw Speed and Rhythmicity are Reduced in 6-OHDA Rats

Since one of the hallmark symptoms of Parkinson’s disease is bradykinesia^35^, we analyzed paw movement speed and timing. Differences in the timing of the two paws were measured by calculating the time between the start of one paw’s pull cycle and the start of the subsequent pull from the opposite paw (see **Figure 4a**). We hypothesized that, in 6-OHDA rats, the intact paw will pull for longer within each cycle relative to the lesioned paw, reflecting possible compensation for the impaired paw. We found that the time to next pull was longer relative to the lesioned paw in the 6-OHDA rats post lesion (**Figure 4b**, paired t-tests with Bonferroni-Holm correction, n = 9. Pre-lesion: t = 0.589, p = 0.572. Post-lesion: t = 3.04, p = 0.0319, Cohen’s d = 0.916). Sham lesioned animals showed no asymmetry in paw timing (**Supplementary Figure 1e**, paired t-tests with Bonferroni-Holm correction, n = 5. Pre-lesion: t = 0.902, p = 0.836. Post-lesion: t = 0.059, p = 0.956).

**Figure 4:**
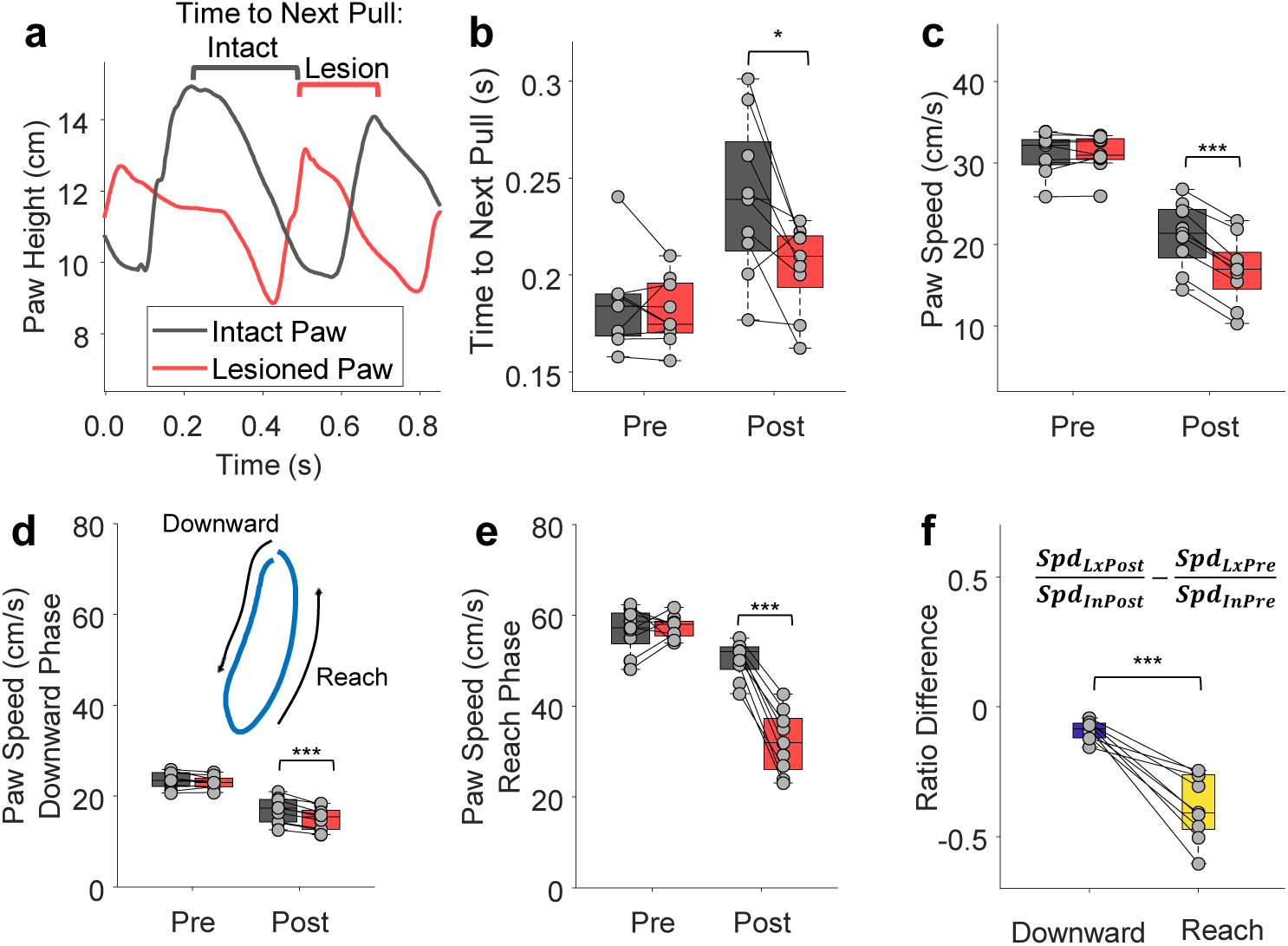
a. Example trace of paw heights during pulling. The time to next pull was defined as the time from the peak of one paw’s pull to the subsequent pull from the opposite paw, shown with a grey bracket for the intact paw and a red for the lesioned paw. b. The time to next pull was significantly longer post-lesion in the intact than in the lesioned paw (paired t-tests with Holm correction, n = 9. Pre-lesion: p = 0.572, post-lesion: p = 0.0319). c. Paw speed was significantly slower in the lesioned paw vs the intact paw post-lesion (paired t-tests with Holm correction, n = 9. Pre-lesion: p = 0.882, Post-lesion: p = 1.22e10-7). d. Paw speed during the reach phase was significantly slower the lesioned paw vs intact paw post-lesion (paired t-tests with Holm correction, n = 9, Pre-lesion: p = 0.203, post-lesion: p = 6.47e-5). e. The change in the ratio of the lesioned paw to the intact paw’s speed pre- vs post-lesion was measured, and we found that the reach phase showed a greater drop than the downward phase (paired t-tests with Holm correction, n = 9. Pre-lesion, p = 0.682. Post-lesion: p = 5.66e-6). ***Spd*** = speed, ***Lx*** = lesioned paw, ***In*** = intact paw, ***Post*** = post-lesion, ***Pre*** = pre-lesion. * = p < 0.05, ** = p < 0.01, *** = p < 0.001

Further, examination of paw speed showed that the lesioned paw was significantly slower than the intact paw following 6-OHDA lesion (**Figure 4c**, paired t-tests with Bonferroni-Holm correction, n = 9. Pre-lesion: t = 0.153, p = 0.882. Post-lesion: t = −14.1, p = 1.22e10-7, Cohen’s d = −4.25), while the sham animals showed no difference (**Supplementary figure 1f**, paired t-tests with Bonferroni-Holm correction, n = 5. Pre-lesion: t = 1.31, p = 0.261. Post-lesion: t = 2.92, p = 0.0864, Cohen’s d = 1.04).

Paw speed changes within each pull cycle with speed peaking during the reach phase^21,22^. Consequently, we independently analyzed paw speed during the reach and downward phases. Given that Parkinson’s disease is associated with impaired effortful movement^11,36^, we hypothesized that paw speed during the downward ‘push and pull’ phase would be particularly impaired post 6-OHDA lesion. Indeed, paw speed during the downward phase was reduced in the intact vs lesioned paw following 6-OHDA lesion (**Figure 4d**, paired t-tests with Bonferroni-Holm correction, n = 9. Pre-lesion: t = −1.39, p = 0.203. Post-lesion: t = −8.34, p = 6.47e-5, Cohen’s d = −2.51) while sham animals showed no difference (**Supplementary Figure 1g**, paired t-tests with Bonferroni-Holm correction, n = 5. Pre-lesion: t = 2.64, p = 0.0665. Post-lesion: t = 3.19, p = 0.0665). However, reach speed was also significantly reduced relative to the intact paw post lesion (**Figure 4e**, paired t-tests with Bonferroni-Holm correction, n = 9. Pre-lesion: t = 0.426, p = 0.682. Post-lesion: t = −11.6, p = 5.66e-6, Cohen’s d = −3.48). Sham animals showed no difference across paws in paw speed during the reach phase (**Supplementary Figure 1h**, paired t-tests with Bonferroni-Holm correction, n = 5. Pre-lesion: t = 1.23, p = 0.435. Post-lesion: t = 1.46, p = 0.435).

We next compared paw speed during the downward and reach phases by first computing the ratio of the paw speeds (lesioned paw speed divided by intact paw speed) before and after 6-OHDA lesion. The pre-lesion ratio was then subtracted from the post-lesion ratio (see **Figure 4f inset**). This analysis revealed that the reach phase showed a greater drop relative to the withdraw phase post lesion (**Figure 4f**, paired t-test, n = 9, t = 6.46, p = 0.000197, Cohen’s d = 1.94). This suggests that, while paw speed is disrupted post lesion, it is the reach phase that is preferentially affected.

### 6-OHDA Lesion Does Not Reduce Effort when Measured as String Tension

Individuals with Parkinson’s disease express difficulties with effortful actions such as sustained grip force^1^. To investigate whether 6-OHDA lesion reduced effort during string pulling, we measured the tension applied to the string during pulling. This was accomplished using a load cell integrated with the pully apparatus (**Figure 5a,b**, see Methods). An example of the readout from the tension sensor is shown in **Figure 5c**. Using a subset of lesioned rats and all sham-lesioned rats we recorded string pulling with the tension sensor. Contrary to our hypothesis, average tension during pulling was not different between groups (**Figure 5d**, unpaired t-test, n = 5 sham, 6 lesion, t = 1.58, p = 0.148). That said, and as shown in **Figure 5d**, one rat in the lesioned group appeared to be an outlier. However, there was nothing unusual about this animal, and it was not excluded from the analysis.

**Figure 5:**
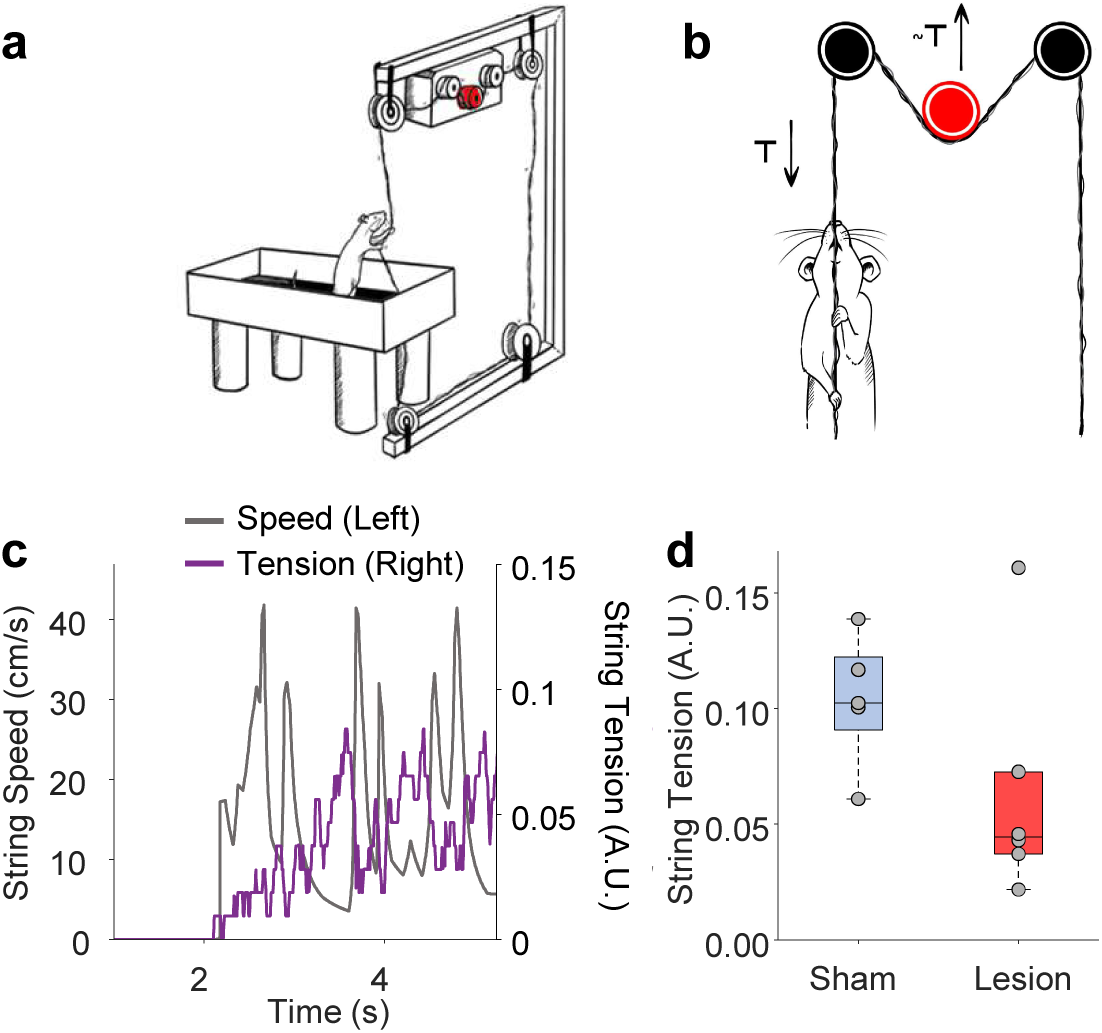
a. String pulling schematic set up showing tension meter in the loop. b. Tension meter measures relative changes in force applied to the string. c. Example trace showing string tension vs string speed. d. Average force applied to string was not significantly different between sham and lesioned animals (unpaired t-test, n = 5 sham-lesion, 6 6OHDA-lesion rats, p = 0.148) * = p < 0.05, ** = p < 0.01, *** = p < 0.001

### L-DOPA Fully Restores Bout Duration and Partially Restores String Speed

Previous work has shown that L-DOPA treatment can restore gross motor skills to near baseline levels^37–40^. Using the same lesioned rats, we measured string pulling after L-DOPA treatment (6 mg/kg L-DOPA with 12 mg/kg Benserazide, i.p.). We excluded 3 rats for showing signs of dyskinesia. Both average string speed (**Figure 6a,b** RMANOVA, F = 24.87, df = 2, n = 8, p = 2.464e-05. Paired t-tests with Bonferroni-Holm correction, n = 8. Pre-lesion vs post lesion: t = 5.94, p = 0.00172, Cohen’s d = 2.7. Post-lesion vs L-DOPA: t = −3.44, p = 0.0109, Cohen’s d = −1.08. Pre-lesion vs L-DOPA: t = −4.97, p = 0.00325, Cohen’s d = −1.56) and peak string speed (**Figure 6c**, RMANOVA, F = 31.62, df = 2, n = 8, p = 6.428e-06. Paired t-tests with Bonferroni-Holm correction, n = 8. Pre-lesion vs post-lesion: t = 8.60, p = 0.00172, Cohen’s d = 1.87. Post-lesion vs L-DOPA: t = −2.89, p = 0.0233, Cohen’s d = −0.908. Pre-lesion vs L-DOPA: t = 5.12, p = 0.00274, Cohen’s d = 1.61) were significantly faster after L-DOPA treatment but were still significantly slower than pre-lesion. Mean bout duration after L-DOPA treatment, however, was significantly increased and was not significantly different from baseline (**Figure 6d**, RMANOVA, F = 12.25, n = 8, df = 2, p = 0.000839. Paired t-tests with Bonferroni-Holm correction, n = 8. Pre-lesion vs post lesion: t = 4.6, p = 0.00500, Cohen’s d = 1.44. Post-lesion vs L-DOPA: t = −5.64, p = 0.00235, Cohen’s d = −1.77. Pre-lesion vs L-DOPA: t = −0.0282, p = 0.978). It appeared to be mostly driven by an increase in longer bouts (**Figure 6e**). An autocorrelagram of string speed indicated that rhythmicity was improved after L-DOPA treatment (**Figure 6f**). L-DOPA treatment had no effect on average string tension during pulling (**Figure 6g**, paired t-test, n = 5, t = −1.24, p = 0.284).

**Figure 6:**
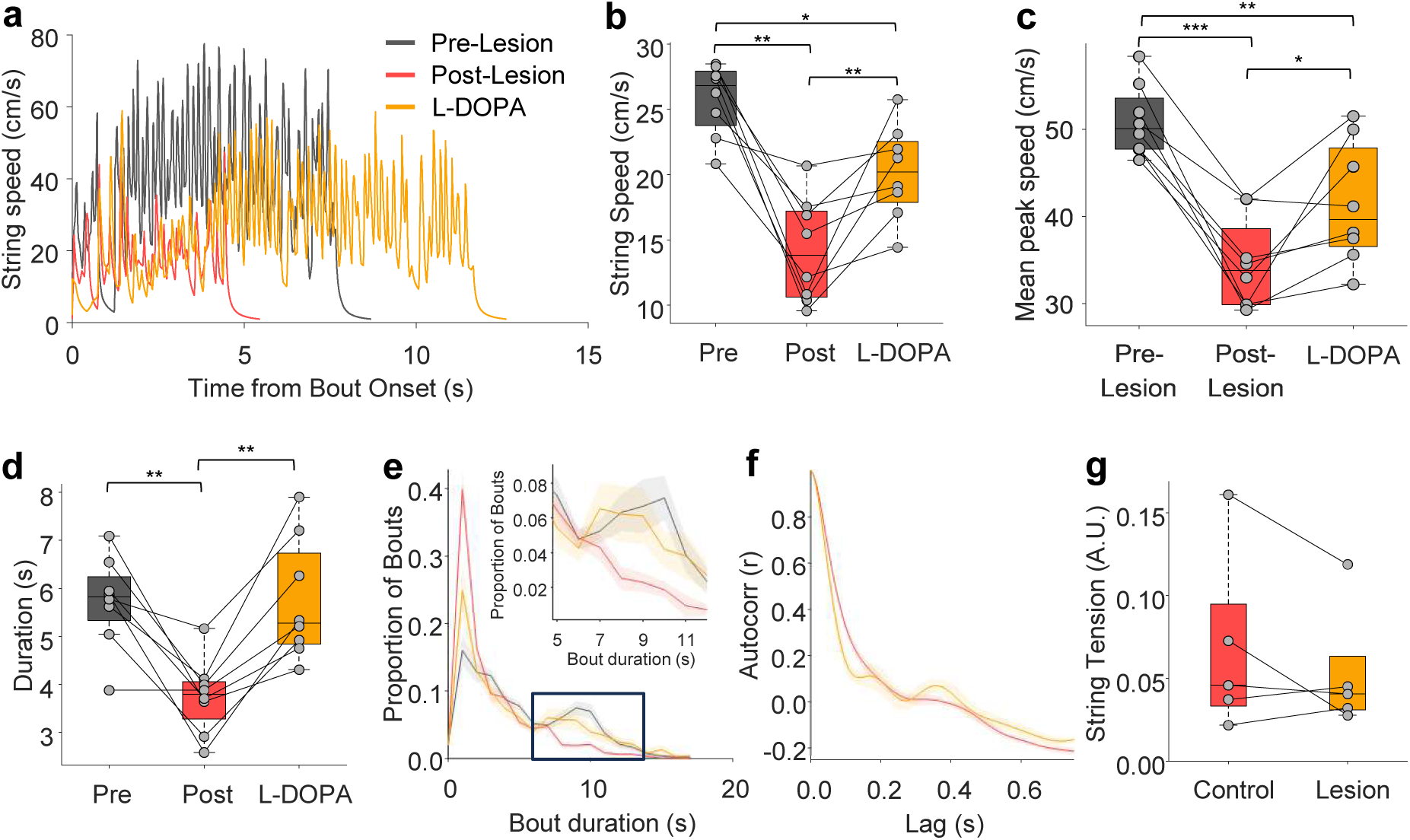
a. Example traces of pulling bouts pre-lesion, post-lesion, and after L-DOPA treatment. b. L-DOPA treatment significantly improved string-pulling speed, but post treatment speed was still significantly different from pre-lesion (RMANOVA, n = 8 rats, p = 2.46e-5, paired t-tests with Holm correction, pre vs post, p = 0.00172, post vs L-DOPA, p = 0.0109, L-DOPA vs pre, p = 0.00324) c. L-DOPA treatment also significantly improved mean peak string-pulling speed, but post treatment peak speed was still significantly different from pre-lesion (RMANOVA, n = 8 rats, p = 6.428e-6, paired t-tests with Holm correction, pre vs post, p = 0.000172, post vs L-DOPA, p = 0.0233, L-DOPA vs pre, p = 0.00274) d. Average bout duration was significantly rescued by L-DOPA treatment; average bout duration after L-DOPA treatment was significantly higher than post-lesion and not significantly different from pre-lesion (RMANOVA, n = 8 rats, p = 0.000839, paired t-tests with Holm correction, pre vs post, p = 0.00499, post vs L-DOPA, p = 0.00235, pre vs LDOPA, p = 0.987). e. e. Average bout duration histograms for pre-lesion, post-lesion, and post L-DOPA, normalized to proportion of all bouts. Inset shows that the proportion of long bouts selectively increases after L-DOPA treatment. Shaded area is SEM across 8 rats. f. Autocorrelogram of string speed post-lesion and after L-DOPA treatment showing improved rhythmicity. Shaded area is SEM across 8 rats. g. g. Average string tension during pulling was not affected by L-DOPA (paired t-test, n = 5, p = 0.284).

### L-DOPA Fully Restores Reach Height but Not Trajectory Area

To determine if L-DOPA restores motor skills and reach/pull movement, we analyzed the paw trajectories of a subset of lesioned rats after L-DOPA injection. The trajectory area was still significantly different across lesioned and intact paws post L-DOPA treatment (**Figure 7a**, RMANOVA, F = 10.62, df = 5, n = 6, p = 1.487e-5. Paired t-tests with Bonferroni-Holm correction, n = 6: Pre-lesion, t = 0.296, p = 0.779. Post-lesion: t = −8.19, p = 0.00132, Cohen’s d = −2.81, L-DOPA, t = −3.84, p = 0.0243). Although the trajectory area was not rescued by L-DOPA, reach height was fully rescued and was no longer significantly different across paws after L-DOPA treatment (**Figure 7b**, RMANOVA, F = 9.36, df = 5, n = 6, p = 0.00265. Paired t-tests with Bonferroni-Holm correction, n = 6. Pre-lesion: t = −1.35, p = 0.470. Post-lesion: t = −12.1, p = 0.000203, Cohen’s d = −4.16. L-DOPA: t = −0.509, p = 0.633). Taken together, these data indicate that L-DOPA preferentially elongated trajectories on the vertical axis.

**Figure 7.**
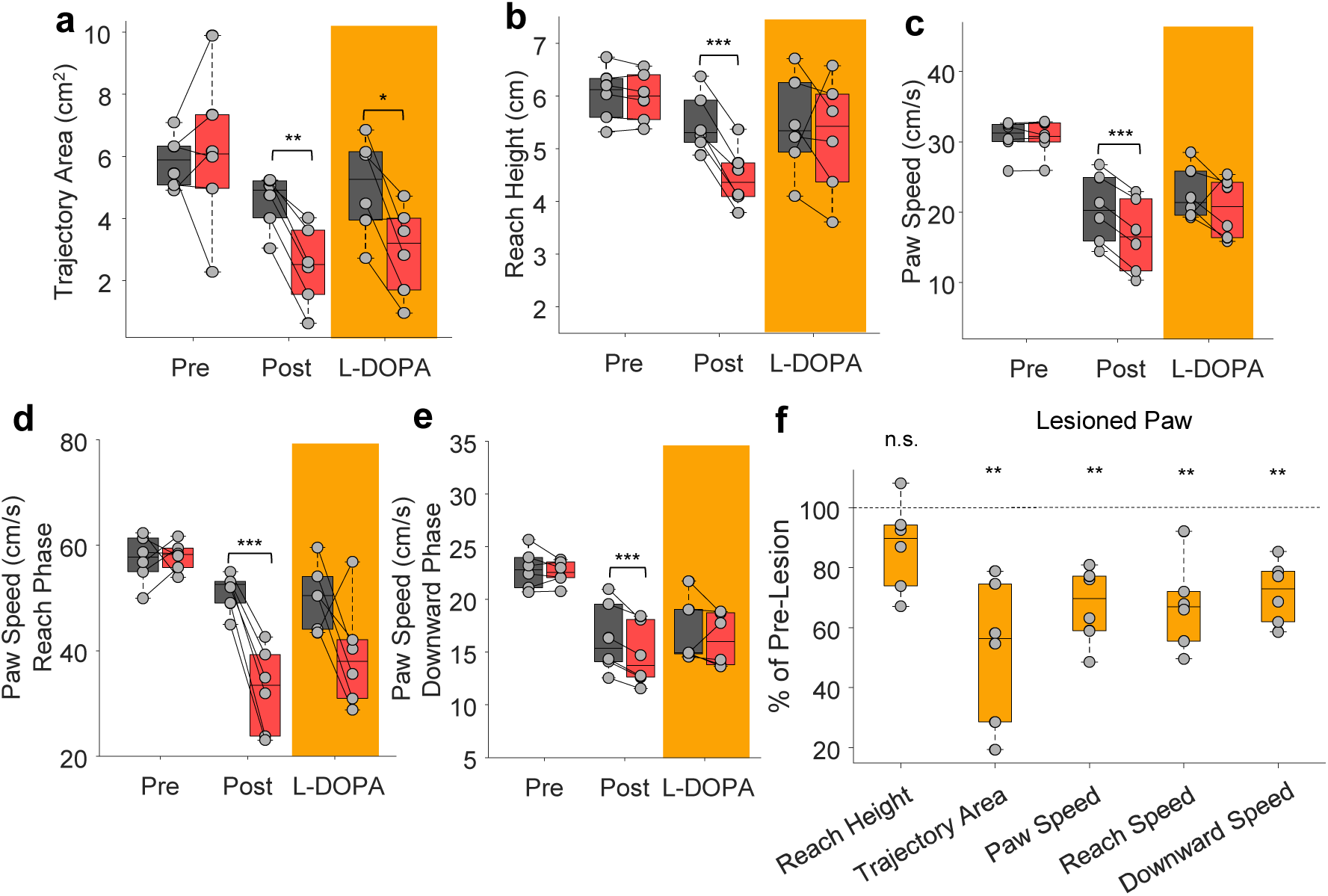
a. Trajectory area was reduced by lesion and was not = restored by L-DOPA treatment (RMANOVA, n = 6 rats, p = 0.00265, paired t-tests with Holm correction, intact vs lesion pre, p = 0.779, post, p = 0.00132, L-DOPA, p = 0.0243). b. Reach height difference lesion vs intact paw was not significantly different post L-DOPA treatment. (RMANOVA, n = 6 rats, p = 1.49e-5, paired t-tests with Holm correction, intact vs lesion pre, p = 0.470, post, p = 2.03e-4, L-DOPA, p = 0.633). c. Post-lesion, lesioned paw speed was significantly different than intact paw speed, post L-DOPA treatment they were not significantly different (RMANOVA, n = 6 rats, p = 0.000330, paired t-tests with Holm correction, intact vs lesion pre, p = 0.661, post, p = 1.09e-5, L-DOPA, p = 0.530). d. Paw speed during the reach phase was no longer significantly different post L-DOPA treatment. (RMANOVA, n = 6 rats, p = 4.99e-08, paired t-tests with Holm correction, intact vs lesion pre, p = 0.824, post, p = 8.19e-4, L-DOPA, p = 0.168). e. Paw speed during the withdraw phase was also no longer significantly different post L-DOPA treatment. (RMANOVA, n = 6 rats, p = 7.144e-11, paired t-tests with Holm correction, intact vs lesion pre, p = 0.979, post, p = 0.00572, L-DOPA, p = 0.979). f. Of all affected movement features, L-DOPA treatment restored only reach height in the lesioned paw to pre-lesion levels (paired t-tests with Holm corrections, n = 6 rats, null hypothesis µ =100%, reach height, p = 0.0876, trajectory area, p = 0.00468, paw speed, p = 0.00148, reach speed, p = 0.00287, withdraw speed, p = 0.00117).

Measuring individual paw velocities, we showed that, after L-DOPA treatment, paw velocity was no longer significantly different across paws, indicating at least a partial rescue (**Figure 7c**). Segmenting paw trajectories into reach and downward phases, we found that reach speed was no longer significantly different after L-DOPA treatment across paws (**Figure 7d**, RMANOVA, F = 20.07, df = 5, n = 6, p = 4.99e-8. Paired t-tests with Bonferroni-Holm correction, n = 6. Pre-lesion: t = 0.234, p = 0.824. Post-lesion: t = −9.07, p = 0.000819, Cohen’s d = −3.11. L-DOPA: n = 5, t = −2.15 p = 0.168), and neither was downward speed (**Figure 7e**, F = 37.79, df = 5, n = 5, p = 7.14e-11, paired t-tests with Bonferroni-Holm correction, n = 6. Pre-lesion, t = −0.746 p = 0.979. Post-lesion: t = −5.96, p = 0.00572, Cohen’s d = −2.04, L-DOPA, n = 6 t = −0.63, p = 0.556).

To compare different movement features with each other, we expressed lesioned paw measures after L-DOPA as percent of pre-lesion levels. Although L-DOPA treatment improved many of our bimanual string pulling measures, only reach height was not statistically different from pre-lesion (Median of 89.77% relative to pre-lesion), suggesting that it was best restored by L-DOPA (**Figure 7f**, RMANOVA, F = 8.00, n = 6, df = 4, p = 5.07e-4. Paired t-tests with Bonferroni-Holm correction, comparing to mean of 100%, n = 6. Reach height: t = −2.12, p = 0.0876. Area: t = −-4.85, p = 0.00936, Cohen’s d = −1.66. Paw speed: t = −6.3, p = 0.00592, Cohen’s d = −2.16. Reach speed: t = −5.43, p = 0.00861, Cohen’s d = −1.86. Withdraw speed: t = −6.64, p = 0.00583, Cohen’s d = −2.28). Together these suggest that L-DOPA can improve many aspects of string pulling but only reach height and bout duration approached pre-lesion levels.

## Discussion

Here we demonstrate that our string-pulling paradigm, with automatic string speed, tension measures, and paw tracking, is a powerful tool for studying how Parkinson’s disease affects motors skills and limb coordination. We showed that unilateral 6-OHDA lesion reduces string speed and pulling bout duration but has no effect on the force applied to the string. We also showed that the paw controlled by the lesioned hemisphere moves slower than the opposite paw post lesion, pulls for less time, and that its movement trajectory becomes smaller. Further, by segmenting movement into reaching and downward phases we found that the paw speed during reaching is most affected by the 6-OHDA lesion. Most other gross motor^15,39,41,42^ and fine motor^9,10,43,44^ tasks provide only a fraction of this information, giving new access to measures of bimanual motor performance in a rodent model of Parkinson’s disease.

The changes we observed here parallel motor deficits in individuals with Parkinson’s disease. For example, paw and string speed reductions reflect bradykinesia seen in Parkinson’s disease^35,45^, the reduced bout duration reflects difficulty executing longer movement sequences^46^, and reduced reach height and trajectory size correspond to reduced movement extent and hypokinesia^47^.

The reduced trajectory area in 6-OHDA rats observed here has striking parallels to micrographia in Parkinson’s disease^30^. As the disease progresses, handwriting becomes smaller and cramped ^30,48,49^. Similarly, all rats showed smaller trajectory area following 6-OHDA lesion. In human literature, stroke speed and acceleration also becomes slower with disease progression^48,50^, similar to our observed reduction in paw speed. String-pulling parallels handwriting in that both are repeatedly practiced and repetitively produced motor tasks. String pulling may be an effective animal model for these kinds of well-practiced skills where Parkinson’s disease alters the movement trajectory.

Other limb motor control tasks in rodents do not provide the same detail and quantitative analysis that the string-pulling task does. The Vermicelli handling task measures time to consume the pasta and number of adjustments, as well as a list of qualitative ways of manipulating the pasta, but does not have any quantitative automated measures of motor performance^9^. Similarly, the staircase test only measures the number of pellets retreived^43^. The cylinder test, an ubiquitous test in the Parkinson’s literature, only measures the relative use of each paw and is rater intensive^42^. There are several tracked lever pressing and pellet retrieval tasks in rodents, but they do not produce the same number of repeated movements, and none require bimanual coordination like the sting-pulling task does. Gait analysis gets the closest to our technique and has the benefit of being highly applicable to the everyday life of people with Parkinson’s disease. Recent progress has been made in this direction in Parkinson’s disease^51,52^. However, the involvement of lower motor nuclei in gait, and the resistance of gait deficits to L-DOPA therapy, suggests the mechanisms may differ^53^.

The string-pulling task is well-suited for the standard 6-OHDA hemi-lesioned model of Parkinson’s disease as it allows comparisons not only between sham and lesioned animals and between the intact and lesioned hemispheres. In addition, the string-pulling behavior partly addresses the challenge of dissociating motivational and motor effects. Parkinson’s patients experience apathy that can affect the effort they exert on a rewarded task in addition to motor symptoms^54,55^ Studies have also shown that rats execute motions that are more rewarded more vigorously^56,57^. However, the unilateral 6-OHDA model provides a strong internal control for motivation in the contralateral, intact paw. Many of our conclusions are based on this post-lesion asymmetry.

Since other tasks assessing limb coordination in animal models do not have the same rich repertoire of measurements, our string-pulling behavior could have a large impact on characterizing the effect of pharmacological treatments on motor control in Parkinson’s disease. In humans, L-DOPA treatment does not restore motor function uniformly. Above we mention how finger tapping, a common diagnostic of Parkinson’s disease, was not significantly improved by L-DOPA treatment ^19^. It has also been shown that stroke size as well as repeated-letter size decline characteristic of micrographia in Parkinson’s disease is not improved by L-DOPA treatment^49,58^. Together these suggest that L-DOPA may be less effective at improving fine-motor than gross-motor skills. Here, in our 6-OHDA lesioned rats, we showed that L-DOPA treatment improved string speed, bout duration, paw velocity and reach height, but did not restore trajectory area difference across paws at all, mirroring how L-DOPA was ineffective at improving micrographia. Coordinated motor skills may be more dependent on dynamic dopamine changes^59,60^, which L-DOPA treatment, producing a 2-3 hour increase in brain dopamine levels in 6-OHDA rats when taken^61^ is unlikely to fully restore.

The string-pulling paradigm gives us detailed access to complex bimanual motor performance relevant to improving human quality of life in an animal model of Parkinson’s disease. By pre-training the rats and measuring performance post-lesion and post-retraining, we are able to elicit a rich repertoire of behavior from a severely lesioned animal. Combined with our automated measurements of task performance and motor behavior, we have access to subtle differences that can be used to more accurately parse changes in limb coordination in future studies using the rat as a model.

### Limitations

A limitation of the present study as that we used a model of late-stage Parkinson’s disease. A future direction is to determine if measures described here can be used to investigate the deficits in models of early-stage Parkinson’s disease and across disease progression in a progressive model. A second limitation is the use of only male rats, and so future work should incorporate female rats to test for any sex specificity. A further limitation is that we only tested after re-training, potentially missing motor deficits associated with early stages of re-training. We are specifically studying a task re-learned after a 6-OHDA lesion. Future studies could explore the early stages of re-training.

Another caveat to consider is the strong link between motivation and movement. Studies have shown that rats execute motions that are more rewarded more vigorously^56,57^, and 6-OHDA lesion of the medial forebrain bundle also lesions dopaminergic projections to the nucleus accumbens, causing motivational deficits in rats^62^. Compounding this, the increased difficulty of movement post lesion also affects the perceived reward of working for food via string pulling^63,64^. However, the unilateral 6-OHDA model again provides astrong internal control for motivation in the contralateral, intact paw. Many of our conclusions are based on this post-lesion asymmetry.

