## Supplementary figure 1. for "L-DOPA Rescues Reach Height but Not Trajectory Area in a Bimanual String-Pulling Behavior in Parkinsonian Rats"

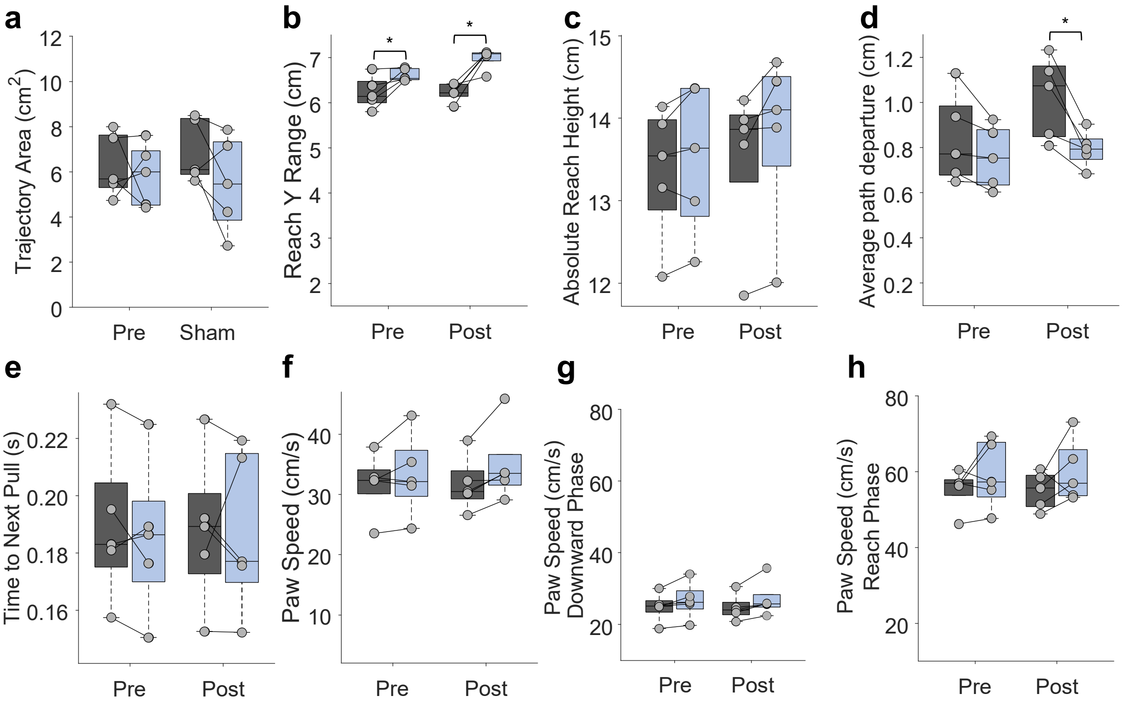


Supplementary Figure 1

a. Trajectory area was not affected post sham-lesion (paired t-tests with Holm correction, n = 5. Pre-lesion: p = 0.669, post-lesion: p = 0.314)
b. Y reach range was significantly higher in the sham lesioned paw both pre and post lesion. (paired t-tests with Holm correction, n = 5. Pre-lesion: p = 0.0392, post-lesion: p = 0.0192).
c. Reach heigh was not significantly different across paws pre- or post-lesion (paired t-tests with Holm correction, n = 5. Pre-lesion: p = 0.189, post-lesion: p = 0.179).
d. Mean path departure in the sham-lesioned paw was significantly smaller post lesion post-lesion (paired t-tests with Holm correction, n = 5. Pre-lesion: p = 0.095, post-lesion p = 0.0335).
e. Time to next pull was not significantly different across paws pre- or post lesion (paired t-tests with Holm correction, n = 5. Pre-lesion: p = 0.902, post-lesion p = 0.956).
f. Paw speed was not significantly different across paws pre- or post lesion (paired t-tests with Holm correction, n = 5. Pre-lesion: p = 0.261, post-lesion p = 0.0864).
g. Downward speed was not significantly different across paws pre- or post lesion (paired t-tests with Holm correction, n = 5. Pre-lesion: p = 0.0655, post-lesion p = 0.0655).
h. Reach speed was not significantly different across paws pre- or post lesion (paired t-tests with Holm correction, n = 5. Pre-lesion: p = 0.435, post-lesion p = 0.435).
